# Quantification and Isolation of *Bacillus subtilis* Spores using Cell Sorting and automated Gating

**DOI:** 10.1101/528257

**Authors:** Marianna Karava, Felix Bracharz, Johannes Kabisch

## Abstract

The Gram-positive bacterium *Bacillus subtilis* is able to form endospores which have a variety of biotechnological applications. Due to this ability, *B. subtilis* is as well a model organism for cellular differentiation processes. Sporulating cultures of *Bacillus subtilis* form sub-populations which include vegetative cells, spore forming cells and spores. In order to readily and rapidly quantify spore formation we employed flow cytometric and fluorescence activated cell sorting techniques in combination with nucleic acid fluorescent staining in order to investigate the distribution of sporulating cultures on a single cell level. Moreover we tested different fluorescent dyes as well as different conditions in order to develop a method for optimal separation of distinct populations during sporulation. Automated gating procedures using k-means clustering and thresholding by gaussian mixture modeling were employed to avoid subjective gating and allow for the simultaneous measurement of controls. We utilized the presented method for monitoring sporulation over time in strains harboring different genome modifications. We identified the different subpopulations formed during sporulation by employing sorting and microscopy. Finally, we employed the technique to show that a double knock-out mutant of the phosphatase gene encoding Spo0E and of the spore killing factor SkfA results in faster spore formation.

## Importance

The sporulation process of the Gram-positive bacterium *Bacillus subtilis* is of broad interest both in basic research as a model for cell differentiation, as well as in biotechnology as tool for displaying heterologous proteins on the spores surface. In the present study we report the development of a high throughput procedure for rapid and robust quantification of *B. subtilis* spores in culture. For the first time, we monitor the formation of *B. subtilis* endospores by flow cytometry and use this method to characterize spore-forming mutants. The described methodology allows for automated, direct characterization of sporulation independently of germination. The method can be applied for absolute quantification of sporulating cells and spores for different time points and mutants.

## Introduction

*Bacillus subtilis* is a model organism extensively studied for its ability to differentiate depending on the growth conditions (1). Sporulation in *B. subtilis* is one of the most thoroughly investigated cellular differentiation programs, triggered by a combination of signals including nutrient exhaustion and cell density (2, 3). The starving cells undergo an asymmetric division, governed by a complex regulatory network which results in the formation of metabolically inactive spores (4, 5). The levels of the master regulator of this process, Spo0A in its phosphorylated form define whether a cell will enter the sporulation pathway (6). However, even under optimal sporulating conditions, only a part of the population undergoes sporulation (7, 8). Several studies report on the heterogeneity of sporulation as the outcome of microenvironmental signals in combination with genetic fluctuations and stochasticity (6, 7, 9).

*Bacillus subtilis* endospores have been investigated for many years in an effort to elucidate their genetic regulation, their biochemical properties and their morphology (4, 10, 11). A set of unique properties such as resilience to environmental assaults, in combination with the potential of the spore coat to be utilized as a scaffold, laid the ground for numerous biotechnological applications. To date, spores have been utilized as platforms for display of a range of biotechnologically interesting peptides, including enzymes and antigens (12, 13).

Throughout the years, methods have been developed for studying different aspects of sporulation in various species including microscopy, microfluidic chips in combination with fluorescence imaging, Raman spectroscopy and Fourier-transform infrared spectroscopy (FTIR) (14–17). However many described techniques either require special instrumentation or are laborious and not automatable. One of the oldest techniques to date for spores quantification in culture is via hemocytometers (18). Despite the low cost, this technique is tedious and not recommended for studies focused on the dynamics of sporulation. Most commonly quality and quantity of sporulation are determined by counting cell forming units after heat shock treatment (19). This method however is an indirect measure since it can only quantify the survival of the heat treatment. Hence, it can be applied only in cases where the spores are still able to germinate. On the other hand, flow cytometry is a useful, rapid and high throughput technique, employed for evaluation of the intrinsic characteristics of cells or particles based on their light scattering properties (20). Flow cytometry has been employed for many years as a method for single cell analysis and separation of eukaryotic cells. However, so far it has remained challenging to distinguish among bacterial species or small size particles (20, 21). Hence, studies reporting on flow cytometry as a tool for discriminating vegetative cells from spores of *Bacillus subtilis* remain scarce. The here presented method uses DNA staining and flow cytometry with automated gating procedures and enables discrimination of three subpopulations of spores.

## Materials and Methods

### Strains and plasmids

Different strains of *Bacillus subtilis KO7 (Bacillus Genetic Stock Center ID 1A1133, derivative strain of Bacillus subtilis PY79*) were utilized as test organisms in the present study. All strains used are listed in Table 1.

**Table 1.**
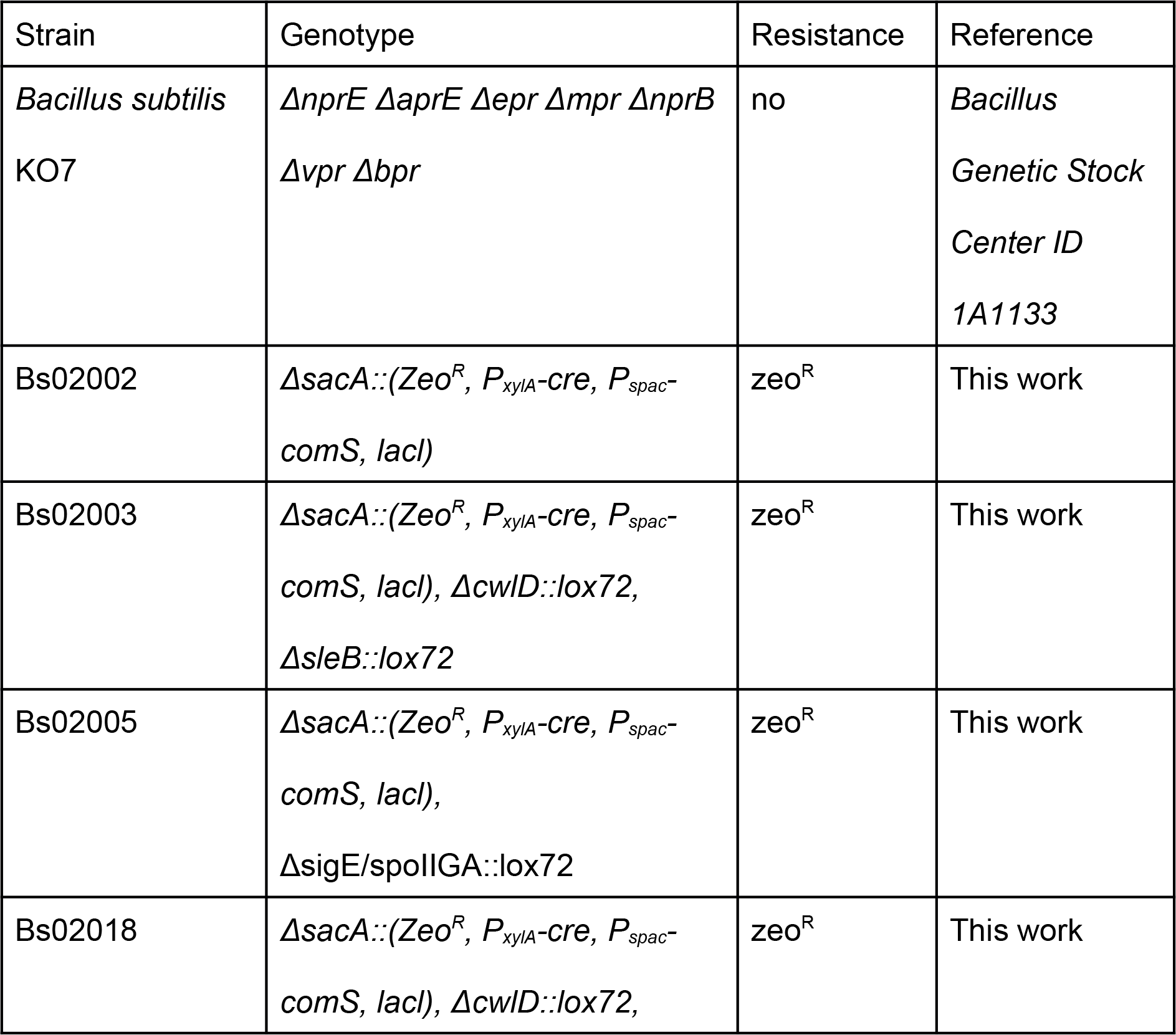
List of strains used in the present study

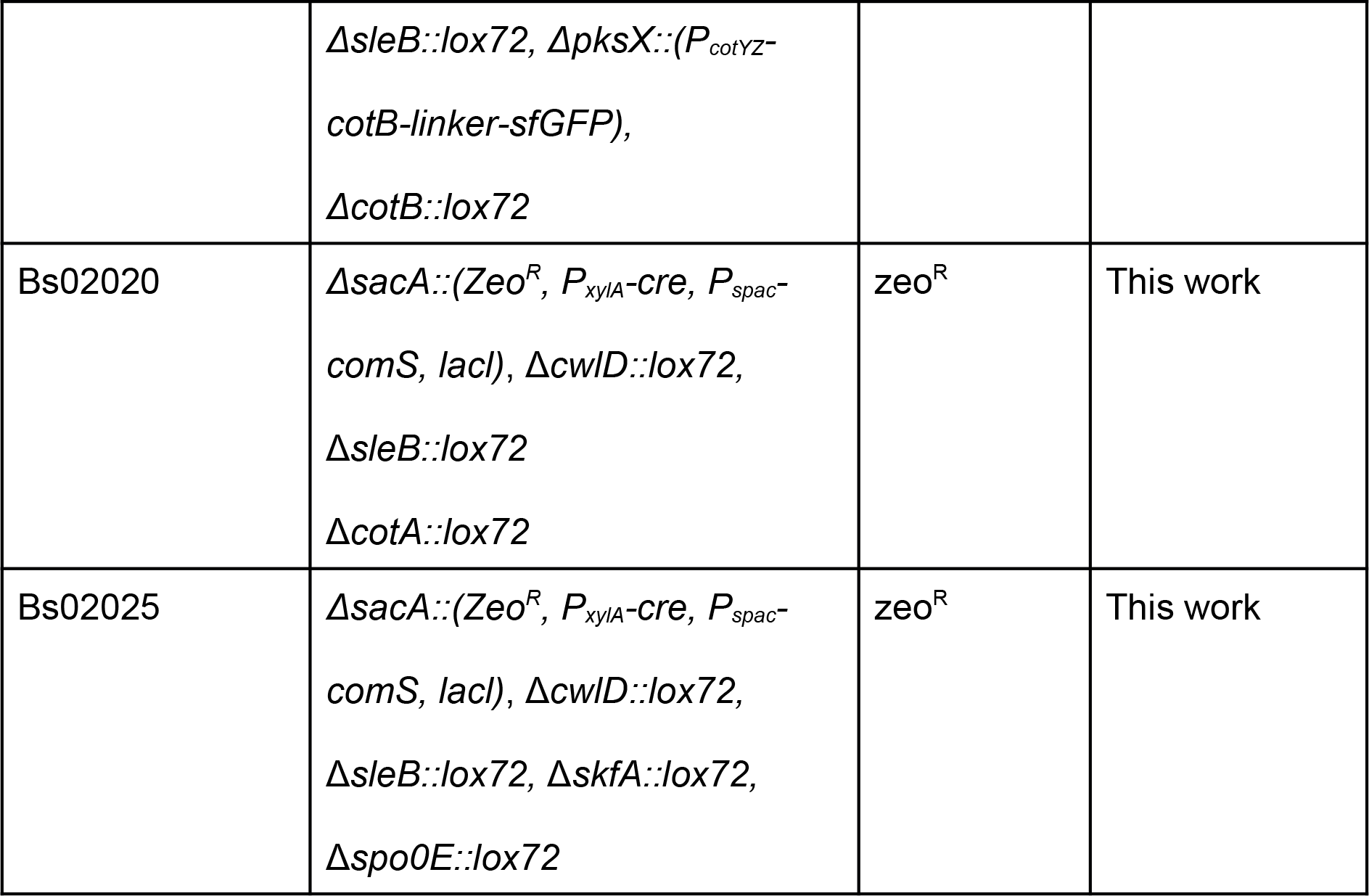

### Media and growth conditions

Vegetative cells were grown in Lysogeny broth (LB) with agitation (200 rpm) at 37°C. Sporulation was induced by nutrient exhaustion as described by Nicholson and Setlow, 1990 (22) including a few modifications. Briefly *Bacillus subtilis* strains were inoculated into 500 ml Erlenmeyer flasks containing 200 ml prewarmed Difco sporulation medium (DSM) and were grown at 37°C with agitation (200 rpm). For spore purification, after 72 h of cultivation the cultures were harvested by centrifugation at 3,400 × g for 15 min. The spores were purified using a Renografin gradient centrifugation as described elsewhere (22) followed by resuspension in 1 × phosphate buffered saline (PBS). The purified spore suspension contained >90% phase bright spores as confirmed by microscopy. For some experimental runs, an artificial mixture of cells and spores with a final optical density at 600 nm of 0.5 was prepared. The artificial mixture contained cells and spores in 1 to 1 proportion.

### Preparation of dyes and staining conditions

Three different fluorescent dyes were utilized for the following experiments: SYBR green I (SGI) (10,000 × stock, Invitrogen™), SYBR Green II (SGII) (10,000 × stock, Invitrogen™) and Propidium iodide (PI) (50 mg, Carl Roth). SGI and SGII stocks were diluted 1:100 times (100 × stock) with class I water respectively. PI working solution was prepared in class I water at 100 mg/ml and employed at a final concentration of 1 mg/ml. For each dye, aliquots of working solutions were transferred in 1.5 ml opaque tubes; aliquots of working dilutions were kept at 4°C. The dyes were tested in different concentrations as well as in combinations. The final tested concentrations of SGI and SGII ranged from 1 × to 4 × of the suppliers suggested concentration while the final concentrations of PI ranged from 0.01 mg/ml to 0.04 mg/ml. Prior to flow cytometry, 500 μl aliquots of cells and spores at an optical density at 600 nm of 0.5 in 1 × PBS, were prepared. After resuspension, the samples were mixed with the appropriate volume of dye and were incubated in room temperature in the dark for 20 min.

### Sample preparation for monitoring sporulation in liquid cell cultures

For each experiment prior to sporulation, precultures were prepared from stock cultures stored at −20°C by streaking the stock solution onto LB agar plates containing zeocin (20 mg/ml). After overnight incubation of the plates at 37°C, a single colony was picked and inoculated into a tube containing 4 ml of LB medium with the respective antibiotic. Cell cultures were grown overnight at 37°C with agitation (200 rpm). Afterwards cells were pelleted by centrifugation at 16,000 × g for 1 min. Supernatant was removed and the cells were resuspended in 2 ml DSM medium. Cell density was defined photometrically at 600 nm and a cell suspension with an optical density at 600 nm of 0.1 was inoculated into 100 ml Erlenmeyer flasks containing 30 ml prewarmed DSM and the respective antibiotic. After 48 hours of inoculation, samples of 200 μl from each culture were acquired for flow cytometry. Samples were centrifuged at 16,000 × g for 1 min with subsequent removal of the supernatant. Pelleted cells were resuspended in 500 μl 1 × PBS and stained with 2x SGI as mentioned above. The same process was repeated for different time points. All experiments were conducted in three biological replicates.

### Flow Cytometry and FACS

Cytometric analyses were done using a Sony LE-SH800SZBCPL with a 488 nm argon laser. Photomultipliers for SSC, FL-1 (525/50 nm) and FL-3 (617/30 nm) were set on 40.0 %, 39.0% and 51.0% respectively with a FSC-threshold of 0.20 % and a window extension of 50. The FSC diode was set on an amplification level of 16/16 and sample pressure was set so that events per second (eps) were kept under 30’000. For analysis and plotting, areas of scatterings- and fluorescence signals were brought to a near-normal form by transforming over the inverse hyperbolic sine.

For cell sorting, the same settings as for cytometric analyses were used, however eps were kept under 12’000. Agglomerates were excluded based on channels FSC-H/FSC-W as well as SSC-H/SSC-W.

### Viability of cells and spores after staining

To monitor the effect of staining in the viability of cells and spores, distinct samples were stained with PI, SGI and SGII as mentioned above. Viability assessment was performed by sorting 210 events from each sample on an LB plate containing zeocin (20 mg/ml). Subsequently, plates were incubated overnight at 37°C. For analysis, the relative number of colonies in respect to the total number of sorted events was evaluated.

### Data analysis and clustering

R package flowcore (23) was used to import fcs files and exclude agglomerates from analysis. Single cells were gated by fitting bivariate normal distributions to channel pairs FSC-W/FSC-H and SSC-W/SSC-H. If not stated differently, a scaling factor of 1 was used. For separation of artificial 1:1 spore:cell mixtures, distributions were separated by assuming a mixture of two univariate normal distributions and fitting using the mixtools package (23, 24). The cutoff values were then set at the respective signal strength between the two maxima, for which the sum of squared predicted counts of signals is at its minimum. To identify clusters of spores, endospores and vegetative cells, k-means was applied on a reference dataset of a biological sample containing all 3 populations, yielding the 3 centers of the population. For this, k-means was repeated 1000 × and the first clustering with the highest SSwithin/SStotal ratio was selected. Events of all samples from the respective strain were then classified according to their euclidean distance to these centers and counted.

### Microscopy

For phase contrast microscopy 6 × 10^6^ events were sorted in 15 ml tubes. Sorted samples were transferred in 20 ml centrifugal concentrators of 30 kDa molecular cutoff and centrifuged at 3,400 × g for 5 min. Microscopy slides were prepared by pipetting 3 μl of the concentrated sample suspensions on glass slides and covering with glass coverslips. Samples were observed with Axio Vert.A1 (Carl Zeiss) inverted microscope equipped with 100 × oil immersion objective and 10 × ocular magnification. Images were captured with AxioCam ICm1 (Carl Zeiss) camera using ZEN lite 2011 software (Carl Zeiss).

## Results

### Separation of cells and spores from different samples

To develop a robust flow cytometric method for separation of *B. subtilis* vegetative cells and spores, separate samples of vegetative cells and purified spores were initially tested. Samples were respectively stained with SGI, SGII and PI as described in materials and methods and subsequently analyzed by flow cytometry.

As shown in Fig. 1A and 1B, spores exhibit higher SSC and somewhat increased FSC than cells, however differences in scattering behaviour alone does not provide clear distinction of the two populations.

**Fig 1:**
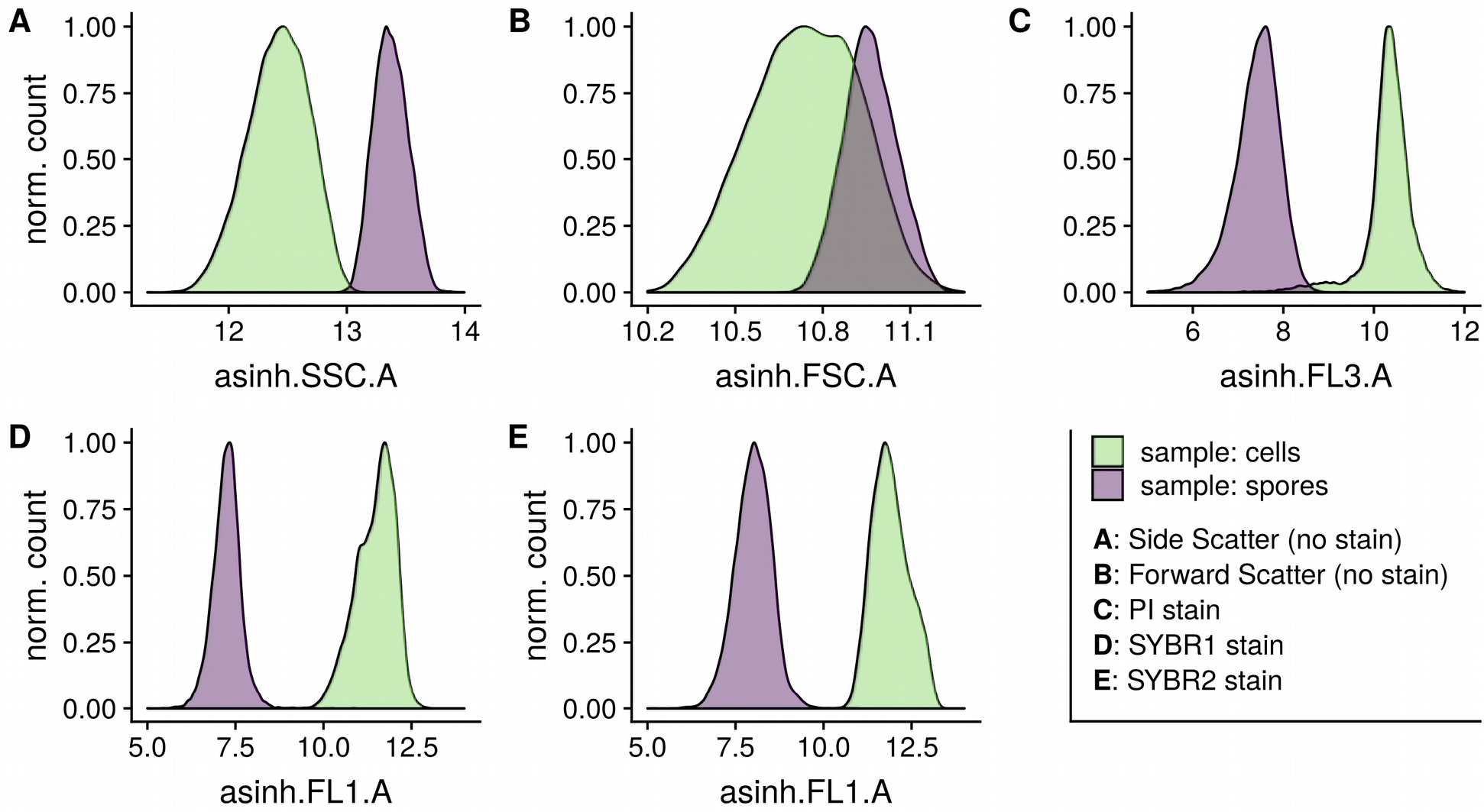
FI and scatter signals of *B. subtilis* vegetative cells and purified spores measured separately. To measure cells the non-sporulating Bs2005 strain was employed. Spores were isolated from a culture of Bs2003 according to Nicholson and Setlow, 1990 (22) resulting in a yield of approx 90%. Characterization of Bacillus subtilis spores with FACS

Staining of samples with fluorescent dyes yielded better separation on the respective channels (Fig. 1C-1E). However, after treatment of cells with PI, partly incomplete staining of cells was observed (Fig. 1C). For SGI and SGII, there appeared to be no such issue (Fig. 1D & 1E).

### Optimization of staining method

To assess the capacity of separating spores from cells coming from the same sample, a 1:1 mix of vegetative cells and purified spores was stained with different concentrations of dye. Subsequently, samples were measured after different incubation times and cell viability was quantified by sorting of cells and spores.

Details of the separation of putative spore and cell populations by gaussian mixture modeling as described in material and methods section. The distance between the distributions of the two populations is shown in Fig. 2 together with the pooled 99.9% confidence interval. Overall, staining with SGI results in the best resolution between cells and spores. Treatment with the two other dyes resulted in smaller differences, whereas minimum difference was observed FSC was used for separation, which is in line with previous experiments.

**Fig 2:**
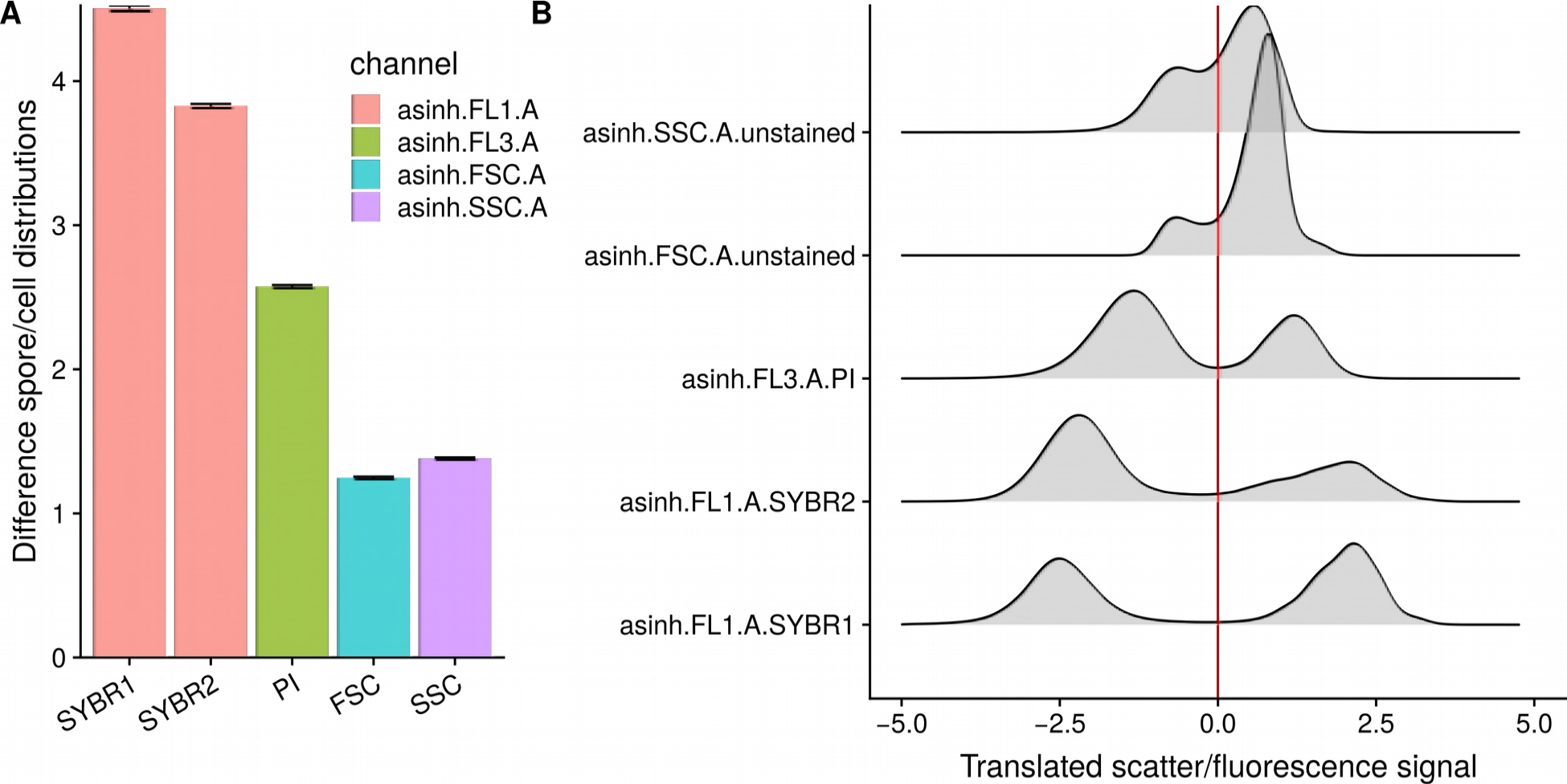
Comparison of different dyes and settings used for optimal separation of cells and spores. A: Distance between population distributions in the respective channel with error bars showing standard deviations. This metric was subsequently used to evaluate the effect of staining time, stain concentration (See supplemental 1: single dye optimization, supplemental 2: mixture optimization) and overall effectiveness of the channel to separate cells from spores. For this, an approximate 1:1 mixture of cells and spores was generated, stained and measured. The 2 main subpopulations were then separated (Supplemental 3). Error bars show 99.9% CIs. 2B: Respective cutoff values based as determined by gaussian mixture modeling. X axis shows translated signal, so that the threshold is always at 0.

Variation of staining time as well as stain concentration yielded only minor differences (Supplemental 1A). Effects of the staining procedure on cell and spore viability was assessed by colony forming units as shown in Supplemental 1B. As a 2-fold concentration after 20 minutes of staining with SGI appears to be sufficient for separation and does not result in significantly increased cell death, this concentration was employed for further investigations.

### Classification and cell sorting of sub-populations

To assess whether the developed method allowed for discrimination of an intermediate stage in the spore formation, sporulating cultures of Bs02003 were tested. Three distinct populations were observed, differing strongly in fluorescence, somewhat in SSC and little in FSC with the centers of the clusters shown in red. To validate that the three discovered subpopulations actually constitute the putative sporulation phases, 6 × 10⁶ events from each cluster were sorted and examined by phase-contrast microscopy (Fig. 3C). Cells found in the respective samples matched the expected pattern of the subpopulation of cells, cells containing endospores and spores.

**Fig 3:**
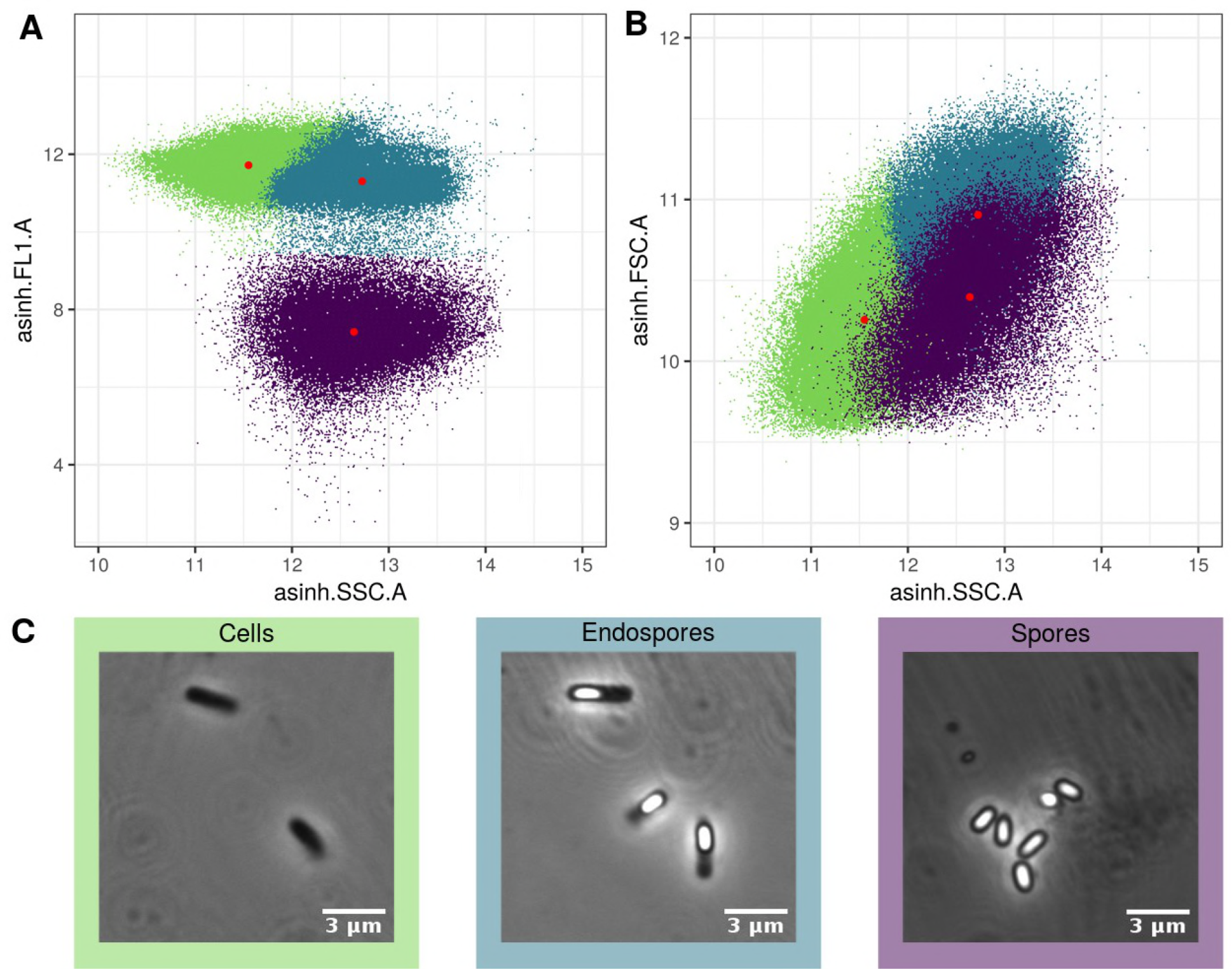
Scatter plots with color indicating the classification. Hyperbolic-sine transformed Side scatter (SSC) and fluorescence in SYBRI channel are shown in A, whereas B shows front- and side scatter (FSC, SSC). Color of the event indicates the respective cluster as predicted by k-means and the respective euclidean distance. Cluster centers are shown in red. For better visibility, translucence indicates log transformed counts. C: corresponding microscopy mages of the respective subpopulations are shown. Subpopulations were isolated by FACS.

### Analysis of sporulation dynamics

The optimized method was applied for quantitative evaluation of sporulation dynamics of *B. subtilis* strains harboring different genome modifications. Initially the sporulation dynamics of strain Bs02018, utilized for display of sfGFP (25) on the surface of spores, was investigated and compared to two other strains harboring similar genomic backgrounds (Bs02003, Bs02020). All three strains carry deletions of both *cwlD* and *sleB*, leading to a germination deficient phenotype (26–28). Strain Bs02020 additionally harbors deletion of *cotA* which is responsible for the brownish pigmentation of spores (29).

For monitoring sporulation of the three strains, population sizes were determined as described above for all strains and time points. In all cases, three distinct populations were observed as shown in Fig. 4A. A clear difference in the sporulation pattern between strain displaying sfGFP (Bs02018) and the two other non-display strains is visible, with Bs02018 exhibiting a delay in sporulation.

**Fig 4:**
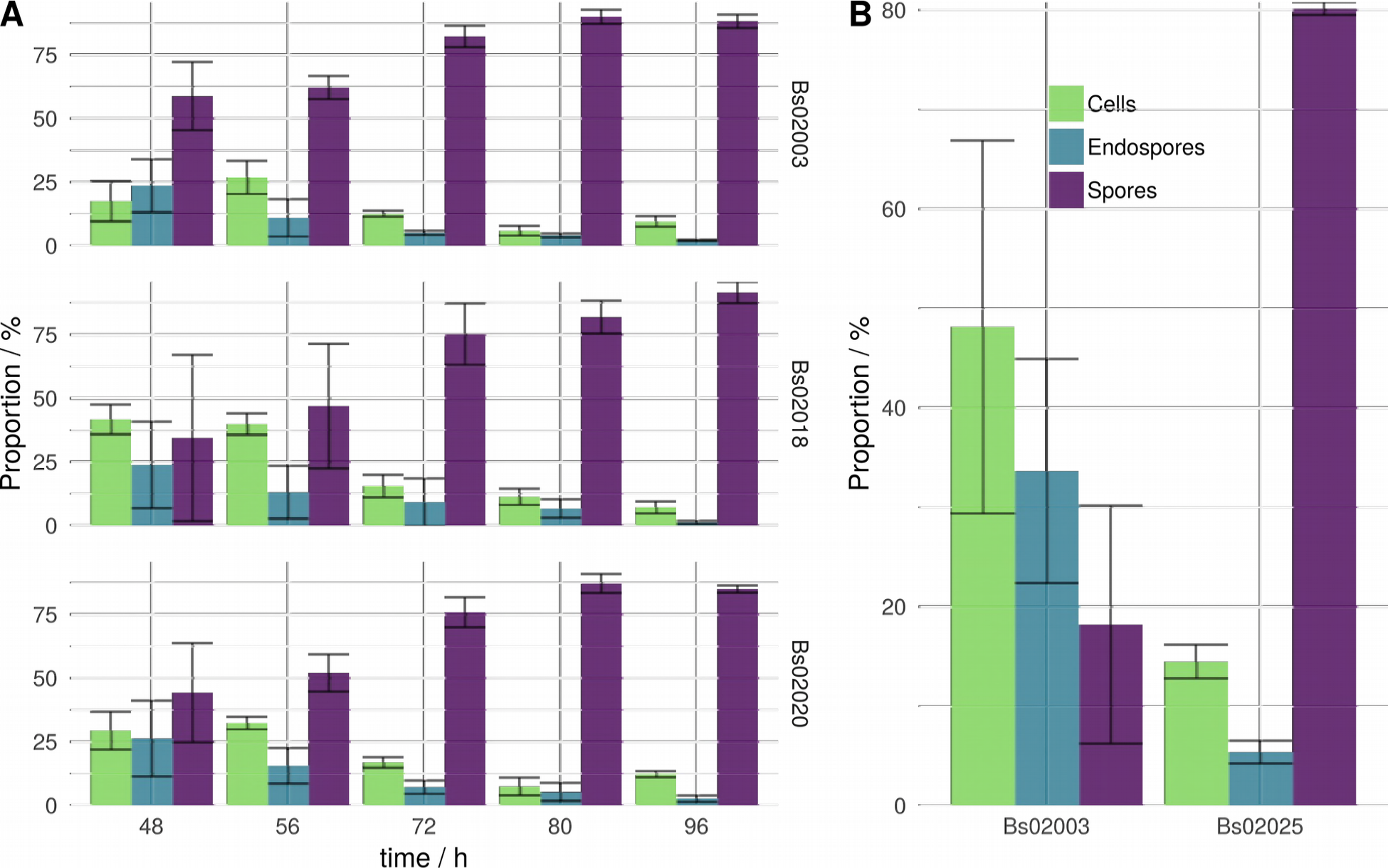
Analysis of sporulation dynamics of *B. subtilis* strains harboring different genome modifications. A: Shift of culture subpopulations containing cells, endospores and spores over time. *S*trains Bs02003, Bs02018 and Bs02020 are double mutants for *cwlD (26)* and *sleB (27)* whereas strain Bs02018 additionally expresses sfGFP as a fusion protein utilized for spore surface display. Strain Bs02020 additionally harbors deletion of *cotA (29)*. 4B: Comparison of culture subpopulations of two *B. subtilis* mutants at 24 hours. Strain Bs02025 contains deletions of *spo0E* (30) and *skfA* (31) whereas Bs02003 serves as control.

As presented in Fig. 4A, already after 48 hours, 59% of events in cultures of strain Bs02003 were spores as opposed to the other strains (35% and 44% respectively). In the course of 96 hours, maximum percentages of spores obtained for the three strains were 88%, 91% and 85% respectively. It is observed that for all strains, only a minority of events was made up by cells without endospores. Furthermore strains Bs02018 and Bs02020 exhibit delayed sporulation in comparison to strain Bs02003. However, in earlier time points high variability was noticed, which made quantification more difficult.

The methodology was further used to evaluate the effect of *spo0E and skfA* deletions. Spo0E is a phosphatase which dephosphorylates the master regulator of sporulation, Spo0A~P and delays the process of sporulation (30) whereas spore killing factor SkfA, is part of the *skf* operon responsible for the production of toxin during sporulation (31). Again, Bs02025 was compared to control strain Bs02003. The generated mutations are associated with highly accelerated sporulation (Fig. 4B). After 24 hours of sporulation, cultures of Bs02025 already consisted of 80% spores while only 18% of events were classified as spores for the control strain.

## Discussion

In this work, we present a flow cytometric method for rapid quantification and isolation of subpopulations formed in sporulating cultures of *B. subtilis*. Spores have more dense internal structure compared to vegetative cells, therefore exhibit increased SSC and somewhat higher FSC. However scattering channels alone are commonly not sufficient for efficient resolution of the two populations. In contrast, Laflamme et al. were able to show separation and isolation of *Bacillus subtilis var niger* spores by employing UV induced autofluorescence (32) with a less common ultraviolet laser. However the necessity for special equipment, hampers the applicability of the described method.

Fluorescent staining has previously been used to investigate the sporulation dynamics of *Paenibacillus polymyxa* and clostridia respectively (33, 34), as well for discrimination of vegetative cells from spores of *Bacillus licheniformis* contained in probiotic tablets (35). In the present study, three different fluorescent dyes were tested for optimal separation of vegetative cells and spores. PI, SGI and SGII are DNA binding dyes commonly used in microscopy and flow cytometry (36–38). The data suggest that SGI is the most suitable dye for separation of subpopulations, whereas PI was the least efficient. This might be attributed to the reduced permeability of vegetative cells for PI. This is in concurrence with the use of PI as a means to separate dead from living cells (39).

Subjective gating and thresholding can lead to different results based on the operators subjective experience (40). When subpopulations can be clearly distinguished, manual methods are mostly sufficient. However, if populations are poorly resolved, inter-lab reproducibility can be achieved better with automated gating (41). Overall interest in automating these processes for flow cytometry has strongly increased over the past years. Thus, we employed different automated methods to allow for more reproducible and operator-independent separation of spore distributions.

In this work, we employed gaussian mixture modeling (GMM) as a means to separate near-normal subpopulations of cells and spores separated on different channels. This allowed for a reduction in tested samples by measuring cell and spore phenotype in a single vial, which further guarantees the same staining environment for cells and spores as opposed to staining separately. GMM is being more routinely used for prediction of subpopulations (42, 43), but is commonly restricted to cases, where signal distributions are at least near-normal. As scattering and fluorescence signals in flow cytometry often follow log-normal distributions, thresholding or clustering with this method can be a useful tool for separation. Further, Lee and Scott (44) showed, that even in the case of censored or restricted data, which frequently occurs in application if the limits of linear detector range is reached, GMM can be modified to suit these conditions.

By using a simple k-means clustering of reference samples of spores, cells and cells containing endospores, the respective subpopulation of any other sample in the respective experiment set was determined by assigning each point to its closest center. If k-means is applied without reference, the same number of populations has to appear in approximately the same place and count, which is mostly not the case in real-life application. In our case, there was little to no shift of populations over the respective independent variable (e.g. time, triplicate). Consequently, euclidean distance to the calculated centers could be used to classify subpopulations. The three detected clusters were attributed to different phases of sporulation based on the following hypothesis: Vegetative cells display high permeability for SGI and exhibit associated high fluorescence intensity. During formation of the endospore, this permeability and FI remain similar, however the emerging protein structure of the coat strongly increase the granularity and hence the measured SSC. Also, a slight increase in average FSC might be attributed to a minor increase in average cell size caused by intracellular formation of the coat. Subsequently, transition from endospore formation to spore release facilitates reduction in FSC and correspondingly cell size, possibly due to smaller size of the spore in comparison to the cell/spore complex. As the spore formation is completed, permeability for SGI decreases due to the presence of the coat or the cortex (45), leading to a drop in FI. Evidence for this hypothesis was gathered by confirming the identity of the respective subpopulations.

It has been demonstrated that the presented method can be utilized for at-line monitoring of sporulation. The optimized method was applied on sporulating cultures of different strains to assess the effect of genomic modifications to the dynamics of sporulation. Quantification of the three major populations formed during sporulation showed a reduction in sporulation efficiency of strain Bs02018, utilized for display of sfGFP on the surface of spores. Based on these results, we assume that expression of sfGFP poses an additional metabolic burden to the already energy consuming process of sporulation. We further demonstrated that a knockout of genes *skfA* and *spo0E* accelerated the sporulation process. These findings are in line with the results reported for *spo0E* deletion (46).

In summary, flow cytometry combined with cell sorting allows for more rapid quantification and isolation of cell populations compared to conventional methods. Our approach facilitates reliable and rapid separation of spores while reducing subjective and laborious gating. We anticipate our method to contribute to the ongoing research in the field of sporulation by allowing for high-throughput analysis using automated incubation and cytometry in microtiter plates as well as for optimizing spore display by enabling rapid characterization of relevant mutants.

## Acknowledgements

MK is supported by the BMBF grant Cascade Kit (FKZ: 031B0579A) and FB is supported by the FNR grant (FKZ: 22007413). JK is supported by a LOEWE CompuGene grant.

The authors have declared no conflicts of interest.

Microbiological work was conducted by MK. Optimization of staining and FACS was done by FB and MK. Analysis and statistical methods were employed by FB. The manuscript was prepared by MK, FB and JK.

## Abbreviations and Symbols

CI: confidence interval
DSM: Difco sporulation medium
Eps: events per second
FL1-A: fluorescence channel 1, signal area (525/50 bandpass) for SGI and SGII
FL3-A: fluorescence channel 3, signal area (617/30 bandpass) for PI
FSC-A: forward scatter, signal area
FTIR: Fourier-transform infrared spectroscopy
LB: Lysogeny broth
PBS: phosphate buffered saline
PI: Propidium Iodide
SGI: SYBR Green 1
SGII: SYBR Green 2
SSC-A: side scatter, signal area

## References

1. Lopez D, Vlamakis H, Kolter R. 2009. Generation of multiple cell types in Bacillus subtilis. FEMS Microbiol Rev 33:152–163.

2. Hilbert DW, Piggot PJ. 2004. Compartmentalization of gene expression during Bacillus subtilis spore formation. Microbiol Mol Biol Rev 68:234–262.

3. Veening J-W, Murray H, Errington J. 2009. A mechanism for cell cycle regulation of sporulation initiation in Bacillus subtilis. Genes Dev 23:1959–1970.

4. Errington J. 1993. Bacillus subtilis sporulation: regulation of gene expression and control of morphogenesis. Microbiol Rev 57:1–33.

5. Piggot PJ, Losick R. 2002. Sporulation genes and intercompartmental regulation, p. 483–517. In Sonenshein, AL, Hoch, JA, Losick, R (eds.), Bacillus subtilis and its closest relatives. American Society of Microbiology.

6. Fujita M, Gonzalez-Pastor JE, Losick R. 2005. High-and low-threshold genes in the Spo0A regulon of Bacillus subtilis. J Bacteriol 187:1357–1368.

7. Chung JD, Stephanopoulos G, Ireton K, Grossman AD. 1994. Gene expression in single cells of Bacillus subtilis: evidence that a threshold mechanism controls the initiation of sporulation. J Bacteriol 176:1977–1984.

8. Kearns DB. 2005. Cell population heterogeneity during growth of Bacillus subtilis. Genes Dev 19:3083–3094.

9. Fujita M. 2005. Evidence that entry into sporulation in Bacillus subtilis is governed by a gradual increase in the level and activity of the master regulator Spo0A. Genes Dev 19:2236–2244.

10. Abhyankar WR, Kamphorst K, Swarge BN, van Veen H, van der Wel NN, Brul S, de Koster CG, de Koning LJ. 2016. The influence of sporulation conditions on the spore coat protein composition of Bacillus subtilis spores. Front Microbiol 7.

11. Plomp M, Carroll AM, Setlow P, Malkin AJ. 2014. Architecture and assembly of the Bacillus subtilis spore coat. PLoS One 9:e108560.

12. Nicholson WL, Munakata N, Horneck G, Melosh HJ, Setlow P. 2000. Resistance of Bacillus endospores to extreme terrestrial and extraterrestrial environments. Microbiol Mol Biol Rev 64:548–572.

13. Isticato R, Ricca E. 2014. Spore surface display. Microbiology Spectrum 2:5.

14. De Gelder J, Scheldeman P, Leus K, Heyndrickx M, Vandenabeele P, Moens L, De Vos P. 2007. Raman spectroscopic study of bacterial endospores. Anal Bioanal Chem 389:2143–2151.

15. Filip Z, Herrmann S, Kubat J. 2004. FT-IR spectroscopic characteristics of differently cultivated Bacillus subtilis. Microbiol Res 159:257–262.

16. Cattoni DI, Fiche J-B, Valeri A, Mignot T, Nöllmann M. 2013. Super-resolution imaging of bacteria in a microfluidics device. PLoS One 8:e76268.

17. Pogliano K, Harry E, Losick R. 1995. Visualization of the subcellular location of sporulation proteins in Bacillus subtilis using immunofluorescence microscopy. Mol Microbiol 18:459–470.

18. Cook AM, Lund BM. 1962. Total counts of bacterial spores using counting slides. J Gen Microbiol 29:97–104.

19. Paidhungat M, Setlow P. 2000. Role of Ger proteins in nutrient and nonnutrient triggering of spore germination in Bacillus subtilis. J Bacteriol 182:2513–2519.

20. Ambriz-Aviña V, Contreras-Garduño JA, Pedraza-Reyes M. 2014. Applications of flow cytometry to characterize bacterial physiological responses. Biomed Res Int 2014:461941.

21. Veal DA, Deere D, Ferrari B, Piper J, Attfield PV. 2000. Fluorescence staining and flow cytometry for monitoring microbial cells. J Immunol Methods 243:191–210.

22. Nicholson W. L. SP. 1990. Sporulation, germination, and outgrowth, p. 391–450. In Harwood, CR, Cutting, SM (eds.), Molecular biological methods for Bacillus. John Wiley and Sons, New York.

23. Hahne F, LeMeur N, Brinkman RR, Ellis B, Haaland P, Sarkar D, Spidlen J, Strain E, Gentleman R. 2009. flowCore: a Bioconductor package for high throughput flow cytometry. BMC Bioinformatics 10:106.

24. Benaglia T, Chauveau D, Hunter DR, Young D. 2009. mixtools: AnRPackage for analyzing finite mixture models. J Stat Softw 32.

25. Lee ME, DeLoache WC, Cervantes B, Dueber JE. 2015. A highly characterized yeast toolkit for modular, multipart assembly. ACS Synth Biol 4:975–986.

26. Sekiguchi J, Akeo K, Yamamoto H, Khasanov FK, Alonso JC, Kuroda A. 1995. Nucleotide sequence and regulation of a new putative cell wall hydrolase gene, cwlD, which affects germination in Bacillus subtilis. J Bacteriol 177:5582–5589.

27. Li Y, Butzin XY, Davis A, Setlow B, Korza G, Üstok FI, Christie G, Setlow P, Hao B. 2013. Activity and regulation of various forms of CwlJ, SleB, and YpeB proteins in degrading cortex peptidoglycan of spores of Bacillus species in vitro and during spore germination. J Bacteriol 195:2530–2540.

28. Setlow B, Melly E, Setlow P. 2001. Properties of spores of Bacillus subtilis blocked at an intermediate stage in spore germination. J Bacteriol 183:4894–4899.

29. Hullo M-F, Moszer I, Danchin A, Martin-Verstraete I. 2001. CotA of Bacillus subtilis is a copper-dependent laccase. J Bacteriol 183:5426–5430.

30. Ohlsen KL, Grimsley JK, Hoch JA. 1994. Deactivation of the sporulation transcription factor Spo0A by the Spo0E protein phosphatase. Proc Natl Acad Sci U S A 91:1756–1760.

31. González-Pastor JE. 2011. Cannibalism: a social behavior in sporulating Bacillus subtilis. FEMS Microbiol Rev 35:415–424.

32. Laflamme C, Verreault D, Ho J, Duchaine C. 2006. Flow cytometry sorting protocol of Bacillus spore using ultraviolet laser and autofluorescence as main sorting criterion. J Fluoresc 16:733–737.

33. Comas-Riu J, Vives-Rego J. 2002. Cytometric monitoring of growth, sporogenesis and spore cell sorting in Paenibacillus polymyxa (formerly Bacillus polymyxa). J Appl Microbiol 92:475–481.

34. Tracy BP, Gaida SM, Papoutsakis ET. 2008. Development and application of flow-cytometric techniques for analyzing and sorting endospore-forming clostridia. Appl Environ Microbiol 74:7497–7506.

35. Zheng X-L, Xiong Z-Q, Wu J-Q. 2017. The use of a simple flow cytometry method for rapid detection of spores in probiotic Bacillus licheniformis-containing tablets. Food Sci Biotechnol 26:167–171.

36. Suzuki T, Fujikura K, Higashiyama T, Takata K. 1997. DNA Staining for Fluorescence and Laser Confocal Microscopy. J Histochem Cytochem 45:49–53.

37. Cerca F, Trigo G, Correia A, Cerca N, Azeredo J, Vilanova M. 2011. SYBR green as a fluorescent probe to evaluate the biofilm physiological state of Staphylococcus epidermidis, using flow cytometry. Can J Microbiol 57:850–856.

38. Riccardi C, Nicoletti I. 2006. Analysis of apoptosis by propidium iodide staining and flow cytometry. Nat Protoc 1:1458–1461.

39. Lehtinen J, Nuutila J, Lilius E-M. 2004. Green fluorescent protein-propidium iodide (GFP-PI) based assay for flow cytometric measurement of bacterial viability. Cytometry A 60:165–172.

40. Aghaeepour N, Finak G, FlowCAP Consortium, DREAM Consortium, Hoos H, Mosmann TR, Brinkman R, Gottardo R, Scheuermann RH. 2013. Critical assessment of automated flow cytometry data analysis techniques. Nat Methods 10:228–238.

41. Burel JG, Qian Y, Lindestam Arlehamn C, Weiskopf D, Zapardiel-Gonzalo J, Taplitz R, Gilman RH, Saito M, de Silva AD, Vijayanand P, Scheuermann RH, Sette A, Peters B. 2017. An integrated workflow to assess technical and biological variability of cell population frequencies in human peripheral blood by flow cytometry. J Immunol 198:1748–1758.

42. Boedigheimer MJ, Ferbas J. 2008. Mixture modeling approach to flow cytometry data. Cytometry A 73A:421–429.

43. Finak G, Bashashati A, Brinkman R, Gottardo R. 2009. Merging mixture components for cell population identification in flow cytometry. Adv Bioinformatics 247646.

44. Lee G, Scott C. 2012. EM algorithms for multivariate Gaussian mixture models with truncated and censored data. Comput Stat Data Anal 56:2816–2829.

45. Magge A, Setlow B, Cowan AE, Setlow P. 2009. Analysis of dye binding by and membrane potential in spores of Bacillus species. J Appl Microbiol 106:814–824.

46. Perego M, Hoch JA. 1991. Negative regulation of Bacillus subtilis sporulation by the spo0E gene product. J Bacteriol 173:2514–252.

